# Machine Learning Approach to Single Cell Transcriptomic Analysis of Sjogren’s Disease Reveals Altered Activation States of B and T Lymphocytes

**DOI:** 10.1101/2024.10.28.620603

**Authors:** Maxwell McDermott, Wenyi Li, Yin-Hu Wang, Rodrigo Lacruz, Bettina Nadorp, Stefan Feske

**Affiliations:** Department of Pathology, New York University Grossman School of Medicine, New York, NY 10016, USA; Department of Molecular Pathobiology, New York University College of Dentistry, New York, NY 10010, USA; Division of Precision Medicine, Department of Medicine, New York University Grossman School of Medicine, New York, NY 10016, USA

## Abstract

Sjogren’s Disease (SjD) is an autoimmune disorder characterized by salivary and lacrimal gland dysfunction and immune cell infiltration leading to gland inflammation and destruction. Although SjD is a common disease, its pathogenesis is not fully understood. In this study, we conducted a single-cell transcriptome analysis of peripheral blood mononuclear cells (PBMC) from patients with SjD and symptomatic non-SjD controls to identify cell types and functional changes involved in SjD pathogenesis. All PBMC populations showed marked differences in gene expression between SjD patients and controls, particularly an increase in interferon (IFN) signaling gene signatures. T and B cells of SjD patients displayed a depletion of ribosomal gene expression and pathways linked to protein translation. SjD patients had increased frequencies of naive B cells, which featured a unique gene expression profile (GEP) distinct from controls and had hallmarks of B cell hyperactivation. Non-negative matrix factorization (NMF) also identified several non-overlapping GEPs in CD4^+^ and CD8^+^ T cells with differential usage in SjD patients and controls. Of these, only the *Th1 activation* GEP was enriched in T cells of SjD patients whereas the other two GEPs were depleted in T cells, emphasizing the important role of Th1 cells in SjD. Our study provides evidence for aberrant and unique gene expression patterns in both B and T lymphocytes of SjD patients that point to their altered activation states and may provide new insights into the pathogenesis of SjD.

## INTRODUCTION

Sjogren’s Disease (SjD) is a systemic autoimmune disease primarily affecting women and the second most common after rheumatoid arthritis^1^. The hallmark characteristic of SjD is the inflammation and dysfunction of salivary glands (SG) and lacrimal gland (LG), which results in dry mouth and dry eyes, also known as Sicca syndrome^2^. The inflammation of exocrine glands is caused by lymphocytic infiltration with T and B lymphocytes and the release of proinflammatory cytokines into the tissues. SjD is characterized by circulating autoantibodies, namely anti-Ro/SS-A and anti-La/SS-B antibodies^3^. While early symptoms and current classification criteria of SjD are predominantly focused on salivary and lacrimal glands, many patients develop extra glandular symptoms due to the involvement of the lungs, kidneys, and other organs in their lifetimes^4^. Patients with SjD also have an increased risk of developing non-Hodgkin lymphoma (NHL) compared to the general population^5,6^. While SjD has not been found to decrease life expectancy, it results in a poor quality of life for those afflicted. Even with this clear clinical need, our understanding of SjD pathogenesis and the corresponding treatment options remain limited.

Currently, a combination of environmental and genetic factors are believed to trigger the onset of SjD^7^. The exact mechanisms underlying SjD pathogenesis remain unidentified, which has also made developing treatment modalities difficult. Most clinical trials have focused on targeting B cells, as not only are they responsible for producing autoantibodies, but they also infiltrate the gland tissues during early disease development^8,9^. The clinical trials targeting B cells have shown promising results in alleviating symptoms of SjD. However, SjD is a complex disorder, with more than one cell type responsible for disease pathogenesis. Both CD4^+^ and CD8^+^ T cells infiltrate the salivary and lacrimal glands of patients with SjD, and in the early stages of disease, they are more abundant than their B cell counterparts^10^. An important role of CD4^+^ T cells in SjD is thought to be the activation and maturation of B cells, but growing evidence indicates that they also play B cell-independent roles in SjD pathogenesis, which are both pro-inflammatory and anti-inflammatory^11–15^. Other cell types besides T and B cells have also been implicated in SjD pathogenesis, including plasmacytoid dendritic cells (pDCs) and monocytes/macrophages^16–19^. pDCs are a specialized subset of DCs that produce large quantities of type I interferon (IFN), and recent studies have noted that expression of the IFN inducible genes is enriched in both SGs and peripheral blood mononuclear cells (PBMCs) of patients with SjD^20^. Similarly, serum interferon levels are elevated in SjD and are currently being researched as both a diagnostic criterion for risk stratification and as a therapeutic target^21^.

In the past, SjD studies have focused on many individual cell types, focusing on their functional status individually^22^. With the advancements in computational biology over the last several years and the emergence of single cell RNA sequencing (scRNA-seq), we can now unbiasedly analyze gene expression in different cell types at the same time in a high throughput fashion with granular precision. In fact, scRNA-seq has been used to study many autoimmune diseases, such as lupus erythematosus (SLE), multiple sclerosis (MS), and rheumatoid arthritis (RA)^23–25^. Recent studies have utilized scRNA-seq to study SjD to find disease-specific cell types and molecular signatures^24,26–32^. As scRNA-seq becomes a common platform to investigate disease pathophysiology, the analysis of such data has become more advanced. Machine learning tools that are advantageous in other fields of science are now being repurposed and applied to scRNA-seq to understand the underlying mechanisms of complex diseases. Deep learning methods are being used to improve several key steps in scRNA-seq analysis, including batch effect removal, cellular clustering, genetic expression imputation and denoising to increase the accuracy of downstream analysis^33^. Non-negative matrix factorization (NMF) uses weighted vectors to model gene expression profiles (GEP) to discover patterns in gene expression in a manner that other methods may not detect. These GEPs can be superimposed onto other datasets, facilitating comparisons across various disease states^34^.

Here, we applied NMF to identify potential mechanisms underlying the pathogenesis of SjD in an unbiased manner. We performed scRNA-seq of PBMCs from 9 patients with SjD and 8 symptomatic controls who did not meet the classification criteria for SjD. NMF was used to assess GEPs in PBMC samples of SjD patients and controls and to detect changes in these GEPs in patients with greater accuracy than standard pathway analyses. PBMCs of patients with SjD were characterized by an enlarged B cell compartment and reduced overall numbers of CD4^+^ T cells. Gene expression was strongly dysregulated in all major immune cell populations of SjD patients, in particular in CD4^+^ and CD8^+^ T cells, B cells, and monocytes. Specific GEPs in CD14^+^ and CD16^+^ monocytes and CD8^+^ T cells were characterized by reduced expression of genes related to the function of these cells. We identified a GEP in B cells that was characterized by increased expression of genes required for B cell function and proliferation, and that was used more heavily by patients with SjD. CD4^+^ T cells were characterized by two distinct functional GEPs, a *Th1 activation* GEP and a *CD4^+^ T cell signal transduction* GEP. The usage of the former was significantly enhanced in SjD patients, whereas the latter was decreased. We show that machine learning approaches are a useful tool to identify unique GEPs associated with altered functional states of immune cells in autoimmune diseases such as SjD. The unique GEPs we identified provide evidence suggesting enhanced B cell and Th1 cell function in patients with SjD.

## RESULTS

### Enrichment of B cells and depletion of CD4^+^ T cells within PBMC of SjD patients

Changes in cellular composition of immune cells within circulating PBMCs and in glandular tissues have been linked to SjD pathogenesis. Whereas most studies have focused on the numbers of immune cell populations, less information is available about their functional states. Here we used single cell transcriptomics data from PBMC of patients with SjD to obtain insights into gene expression and thus, indirectly the function, of different immune cell populations **(Figure 1A)**. With one exception, all nine SjD patients analyzed were positive for anti-SSA/Ro antibodies and had ACR/EULAR scores ranging from 5-9 **(Table S1)**. The control cohort contained eight patients with sialadenitis and sicca symptoms but without SjD (anti-SSA/Ro negative; ACR/EULAR scores of 0 or 1). A total of 196k cells from 17 samples total and three experiments were assembled. Following dimensional reduction and clustering, seven cell types were annotated manually, using conserved marker genes for each major subpopulation **(Figure 1B**), which were validated by reference-based^35,36^ annotation in more detail **(Figure S1A)**. The seven identified main immune cell populations included CD4^+^ and CD8^+^ T cells, other T cell subsets, B cells, NK cells, monocytes and dendritic cells (DCs). Of these, only the frequencies of CD4^+^ T cell and B cell populations showed statistically significant quantitative differences between SjD patients and the control group **(Figure 1C)**. Whereas the frequency of B cells was increased in the SjD patient group, that of CD4^+^ T cells was reduced. An increase in B cell numbers in the blood of SjD patients has been reported^37^, as have reduced numbers of CD4^+^ T cells ^38,39^. Likewise, alterations in the ratio of B and T cells has been demonstrated in patients with SjD although not using scRNA-seq for analysis^40^.

**Figure 1.**
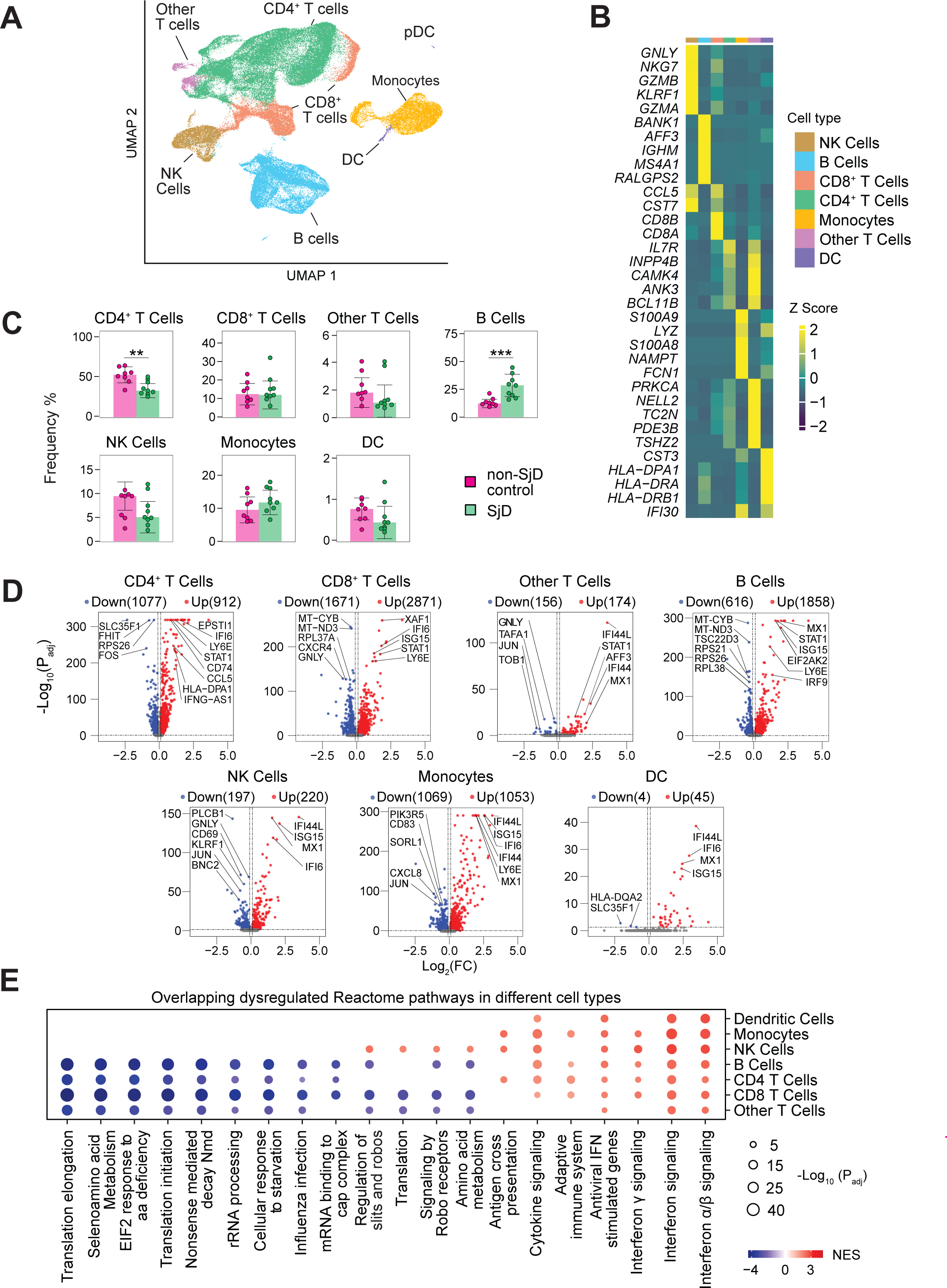
Dysregulated gene expression in immune cell populations within PBMC of SjD patients. **A)** Aggregate UMAP plot of immune cell subsets identified in 105k peripheral blood mononuclear cells (PBMC) from 9 SjD patients and 8 non-SjD controls analyzed across 3 batches. **B)** Heatmap of conserved canonical markers genes of 7 distinct major immune cell subsets. **C)** Median frequency of each major immune cell subset in SjD patients and non-SjD controls. Each dot represents the cell type frequency of one patient. **D)** Volcano plots of differentially expressed genes (DEGs) in SjD patients and non-SjD controls for each major immune cell subset. The top 5 dysregulated genes are annotated. **E)** Dotplot of major dysregulated pathways in each major immune cell subset analyzed by gene set enrichment using genesets from the Reactome database. Shown are overlapping pathways that are dysregulated in at least 3 of 7 cell types. Z scores in (B) were calculated using the normalized average expression per cell in each cell type across all samples. Statistical analyses in (C), (D), and (E) were performed using Wilcoxon rank sum test. Genes in (D) were considered significant if the adjusted P value (P_adj_) was < 0.05. Pathways in (E) were considered overlapping if they were found to have a P_adj_ < 0.01 in 3 or more cell types. Significance was adjusted using the Benjamini-Hochberg method. *P < 0.05; **P < 0.01; ***P < 0.001.

### Dysregulated gene expression in all immune cell populations of SjD patients

We next analyzed if changes in B cell and CD4^+^ T cell frequencies are related to gene expression that might provide clues about their activation state. We analyzed differentially expressed genes (DEGs) between SjD and control samples for each of the main immune cell populations individually using the MAST function of Seurat (**Figure 1D**)^41^. We found strongly dysregulated gene expression in each of the seven immune cell types between SjD and control samples with approximately similar numbers of up- or downregulated genes. The largest numbers of DEGs were found in CD8^+^ T cells, followed by CD4^+^ T cells, B cells, and monocytes. Next, we performed a functional enrichment test using the gene set enrichment analysis (GSEA) function of ClusterprofileR^42^. We found a significant increase in interferon signaling in all seven immune cell populations of SjD patients, with both IFN-γ and IFN-α/β signaling pathways enriched **(Figure 1E)**. Increased type I and II IFN signatures have been reported and are a common molecular feature of SjD^43^. We also observed a strong depletion of pathways related to translational initiation and elongation, rRNA processing, and nonsense-mediated mRNA decay. Depletion of these pathways was restricted to B cells and the three T cell populations and not present in other PBMC subsets. Of note, this analysis focused on shared pathways that are dysregulated in at least 3 of the seven main immune cell populations, whereas differentially regulated pathways in individual immune cell populations of SjD patients will be reported further below. Our data indicate that gene expression is markedly altered in all immune cells within PBMC of patients with SjD and that shared dysregulated pathways mainly affect IFN signaling and ribosome-associated pathways.

### Enhanced intercellular communication of DCs with other immune cells in patients with SjD

Having found that gene expression is markedly altered in all seven immune cell populations of SjD patients’ PBMCs, we hypothesized that these changes may affect intercellular communication. We, therefore, calculated inferred cellular communication between immune cell populations of SjD patients and controls using the Cellchat program^44^. Cell-to-cell communication was analyzed in a weighted manner to accommodate different cell frequencies. Comparing intercellular communication in samples of SjD patients and controls, we found that communication between all cell populations was reduced in samples of SjD patients **(Figure S1B)**. B cells, in particular, showed decreased cellular communication with all other immune cell populations in SjD patients despite their increased frequencies based on scRNA-seq data **(Figure 1C)**. The only cell type that showed increased communication with other PBMC subsets, including all three T cell populations and NK cells, were DCs. We hypothesized that the increased intercellular communication of DCs from SjD patients might correlate with an increase in gene expression compared to controls. A differential gene expression analysis between DCs from SjD patients and non-SjD controls using the Seurat FindMarkers function, however, showed only a small number of genes with increased expression, especially compared to other immune cell types, with less than 50 enriched genes **(Figure S1C)**. We noted a strong upregulation of genes related to IFN signaling in SjD patients. This included MX1 and MX2 and many IFN-stimulated and IFN-inducible genes.

While we were able to identify DEGs in DCs of SjD patients and controls, it is noteworthy that the frequency of DCs in PBMC samples is very small (< 1%) and the number of DEGs much lower than in other immune cell populations (< 50). To avoid missing functional changes based on differential gene expression alone and to better understand the functional status of DCs in patients with SjD, we used non-negative matrix factorization (NMF), a machine learning approach suitable to identify differences between datasets of limited size in an unbiased manner ^45^. NMF allowed us to identify and quantify the functional state of each cell individually while still sequestering the robust IFN signature identified by GSEA. Using NMF, we identified two distinct gene expression profiles (GEP) utilized by the DC population. The first GEP was predominantly used by cells in the pDC compartment, and the second GEP was utilized by all myeloid cells (**Figure 2A, 2B**). When we calculated the usage scores for the first GEP, we found that it was not significantly enriched in pDCs (or conventional DCs) of patients compared to controls, although there was a trend towards preferential usage in the SjD patients (**Figure 2A**). This conclusion, however, is limited by the very small frequency of circulating pDCs. The usage of the second, *All myeloid cell* GEP by DCs was not different between patients and controls either. We did however find a significant, yet small, increase in the usage of this GEP by monocytes of SjD patients (**Figure 2B**). We conclude that although DCs of SjD patients show a marked increase in expression of IFN-stimulated genes, their usage of an *All myeloid cell* GEP is not different from that of non-SjD controls.

**Figure 2.**
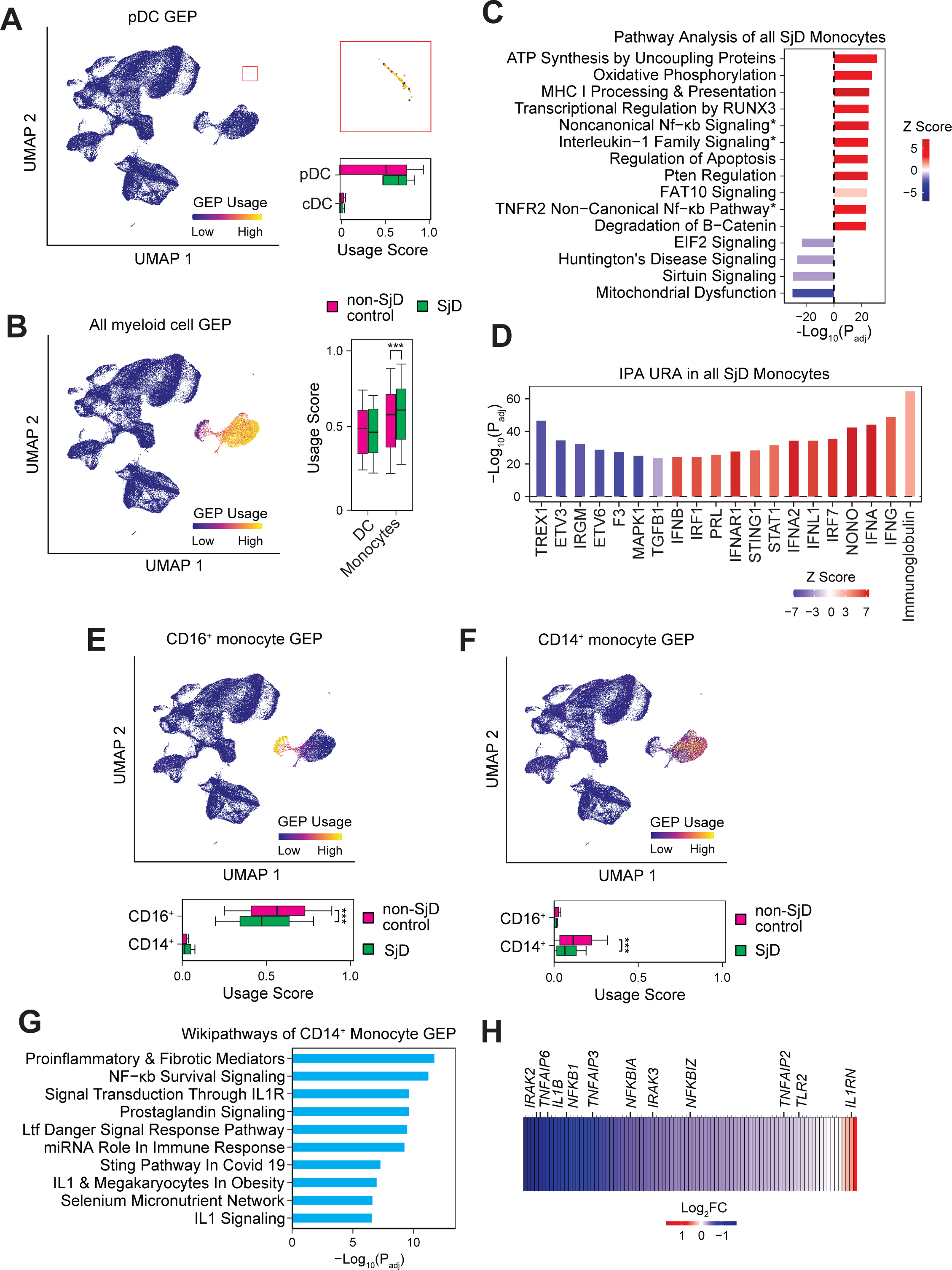
Identification of functional gene expression profiles in dendritic cells (DC) and monocytes of SjD patients. **A)** UMAP plot of PBMCs expressing a gene expression profile (GEP) associated with plasmacytoid dendritic cells (pDC) and identified by NMF. The red box on the right is a zoomed in version of the pDC population shown in the UMAP plot on the left. The bar plot shows a quantification of the usage scores for this GEP in pDCs and conventional DCs (cDCs) in SjD patients and non-SjD controls. **B)** UMAP plot of PBMCs colored by expression of the GEP associated with myeloid lineage cells identified by NMF. The bar plot shows a quantification of the usage scores for this GEP in all DCs and all monocytes in SjD patients and non-SjD controls. **C)** Top 15 dysregulated pathways based on DEGs in all monocytes of SjD patients and non-SjD controls analyzing the canonical pathways of the IPA platform. **D)** Top 20 upstream regulators of DEGs in all monocytes of SjD patients and non-SjD controls using the IPA regulator database. **E)** UMAP plot of PBMCs colored by expression of the GEP associated with CD16^+^ monocytes identified by NMF. The bar plot shows usage scores for this GEP in CD14^+^ and CD16^+^ monocytes in SjD patients and non-SjD controls. **F)** UMAP plot of PBMCs colored by expression of the GEP associated with CD14^+^ monocytes identified by NMF. The bar plot shows usage scores for this GEP in CD14^+^ and CD16^+^ monocytes in SjD patients and non-SjD controls. **G)** Top 10 Wikipathways pathways associated with the CD14^+^ monocyte GEP identified by EnrichR analysis. **H)** Heatmap of DEGs contained in the CD14^+^ monocyte GEP in CD14^+^ monocytes of SjD patients and non-SjD controls. Colors depict log_2_ fold changes (FC) in expression. Statistical analyses in (A), (B), (E), and (F) were completed using Wilcoxon rank sum test. The significance of pathways and regulators shown in (C) and (D) was calculated using a right-tailed Fisher’s exact test. Significance in (A), (B), (C), (D), (E), (F), and (G) was adjusted using the Benjamini-Hochberg method. Scales in (A), (B), (E), and (F) represent the usage scores per cell for each GEP with a range between 0.2 (low) and 0.7 (high). *P< 0.05; **P< 0.01; ***P< 0.001.

### Depletion of distinct functional gene expression profiles in CD14*^+^* and CD16^+^ monocytes of SjD patients

Monocytes and macrophages have not been the focus of much research in SjD, but they may play a significant role in causing salivary and lacrimal gland inflammation through cytokine production, antigen presentation, or phagocytosis^46^. The analysis of transcriptomes of monocytes from SjD patients compared to controls showed the dysregulated expression of more than 2,000 genes **(Figure 1D)**. An IPA pathway analysis of DEGs in monocytes identified several pathways that are enriched in monocytes of SjD patients. These include ATP synthesis and oxidative phosphorylation (with a concomitant depletion of mitochondrial dysfunction-associated genes), MHC class I processing and presentation, IL-1 family signaling, and noncanonical NFκB signaling (**Figure 2C**). Enrichment of the latter two pathways was driven, however, by the presence of many *PSM* genes encoding proteasome subunits in both pathways and not bona fide IL-1 or NFκB signaling genes (not shown). An IPA upstream regulator analysis revealed IFN-related signaling as the main cause of differential gene expression in monocytes of SjD patients (**Figure 2D**). Monocytes could be further segregated into two populations that are distinguished by the expression of CD14 **(Figure S2A)** and CD16 **(Figure S2B)** marker genes. These populations largely overlapped with two functional GEPs that we identified by NMF analysis **(Figure 2E, F)**. The GEP associated with CD16^+^ cells was only used by this monocyte subset but not by CD14^+^ monocytes. The usage of this GEP was reduced in CD16^+^ monocytes from patients with SjD (**Figure 2E**). Conversely, the GEP associated with CD14^+^ monocytes was only used by this monocyte subset. Its usage was also reduced in SjD patients compared to controls (**Figure 2F**).

When we performed a functional enrichment test of genes in the CD16^+^ monocyte-associated GEP, we were unable to identify specific pathways that were dysregulated in SjD patient samples compared to controls, nor were we able to identify distinct patterns of altered genes upon manual inspection **(Figure S2C)**. Although the usage of the *CD16^+^ monocyte* GEP is depleted in SjD patients, the biological relevance of this depletion is unclear. By contrast, a functional enrichment test of the *CD14^+^ monocyte* GEP, which is also depleted in SjD patients, revealed that this GEP is linked to IL-1 and NFκB signaling (**Figure 2G**). Genes belonging to this GEP and depleted in SjD patients include the IL-1 signaling-associated genes *IL1B* (encoding IL-1β), *IRAK2,* and *IRAK3* (encoding interleukin-1 receptor-associated kinases 2 and 3), *TNFAIP3* (encoding the IL-1 signaling inhibitor A20), *IL1RN* (encoding the interleukin-1 receptor antagonist, IRAP), and the NFκB signaling associated genes *NFKB1* (encoding the NFκB p50 subunit), *NFKBIA* (encoding IκBα), and *NFKBIZ* (encoding IκBζ, which is induced by IL-1 signaling). Although NFκB signaling-related genes in the *CD14^+^ monocyte* GEP were depleted, these genes encompassed both positive and negative regulators of NFκB signaling, and the functional consequences of their altered expression remain to be elucidated. It is noteworthy that IL-1 signaling has previously been linked to SjD pathogenesis by controlling systemic or local immune responses and regulating the function of gland epithelial cells^47^.

### Increased abundance and functional signature of naive B cells in patients with SjD

B cells play an important role in SjD by producing autoantibodies, particularly anti-SSA and anti-SSB antibodies. Having determined that the B cell numbers are increased in SjD patients compared to non-SjD controls, we hypothesized that this increase could be restricted to a specific B cell subset. Among the naive, intermediate, and memory B cell subsets identified by scRNA-Seq **(Figure S1A)**, only the naïve subset was significantly expanded in SjD patients, although all subsets showed a trend towards enrichment (**Figure 3A**). Plasmablasts, although detectable, were too infrequent to allow for meaningful biological conclusions; therefore, they were not considered for further downstream analysis. Differential gene expression analysis had shown profound up- and downregulation of a total of 2,774 genes in all B cell subsets combined. A pathway analysis of these DEGs showed enrichment of pathways such as pre-mRNA processing, protein ubiquitination, ATP synthesis, oxidative phosphorylation, non-canonical NF-kB signaling, and BCR signaling, and depletion of pathways including eIF2 signaling, which is crucial for the regulation of protein synthesis (**Figure 3B**). To better understand the causes of dysregulation of such a large number of genes in B cells of SjD patients, we conducted an upstream regulator analysis of DEGs in all B cells of SjD patients and controls using the IPA platform (**Figure 3C**). We found enrichment of several interferons, including IFN-α and IFN-γ as upstream regulators in B cells of patients with SjD. Also enriched was La Ribonucleoprotein 1 (LARP1), which regulates the translation of 5’TOP mRNAs, including those encoding ribosome proteins. Non-phosphorylated LARP1 interacts with 5’ and 3’ UTRs of ribosome-encoding mRNAs and inhibits their translation^48,49^, which is consistent with the decreased activity of pathways associated with translational initiation and elongation observed in B cells **(Figure 1E)**. Depleted upstream regulators in B cells of SjD patients were the proto-oncogenes MYC and MYCN, which is also consistent with the dysregulation of ribosome-associated pathways as MYC regulates ribosome biogenesis and protein synthesis.

**Figure 3.**
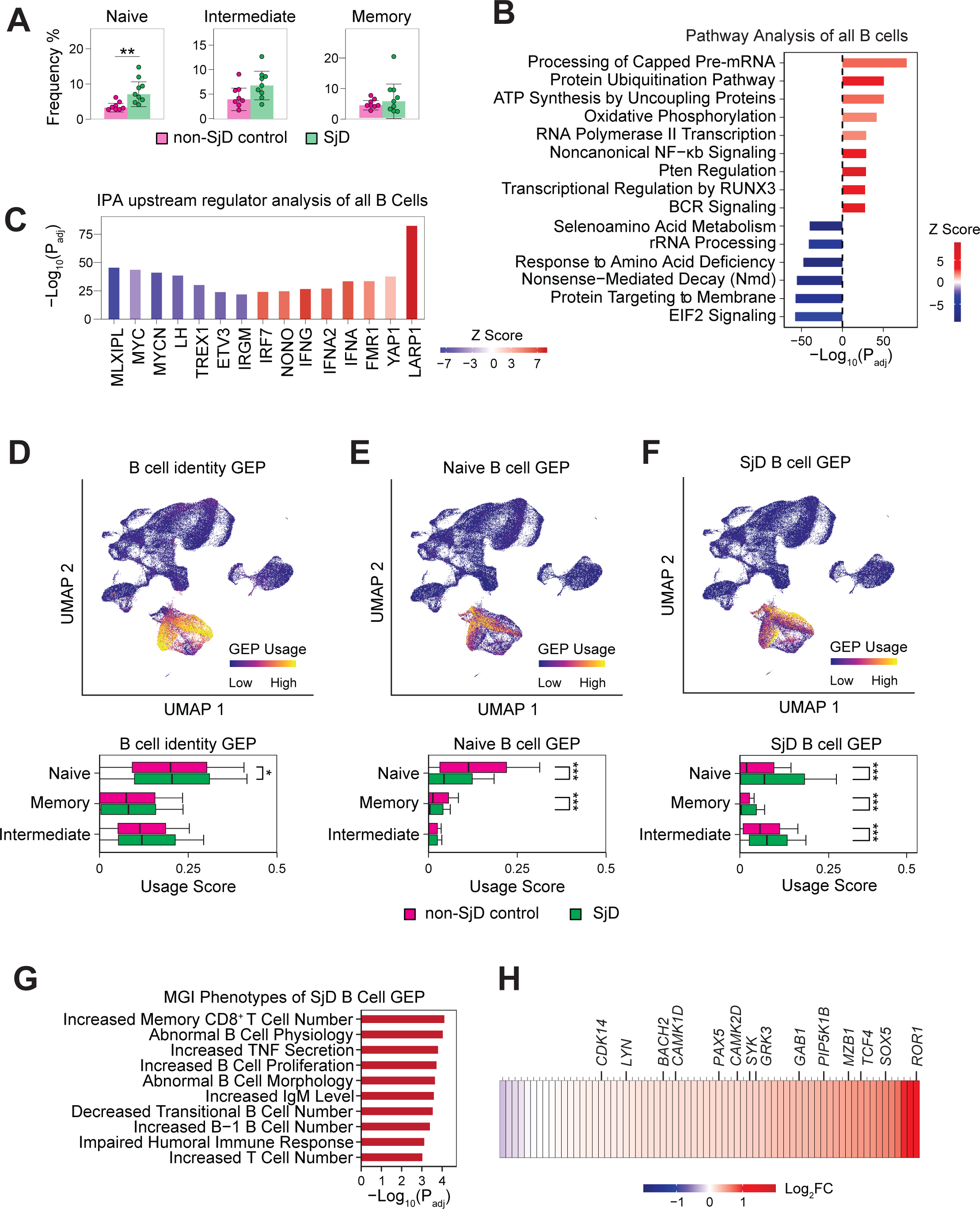
Enrichment of a disease specific GEP in B cells of patients with SjD. **A)** Median frequencies of B cell subsets (of all PBMCs) in SjD patients and non-SjD controls. Each dot represents the cell type frequency of one patient. **B)** Top 15 dysregulated pathways based on DEGs in all B cells of SjD patients and non-SjD controls using the canonical pathways of the IPA platform. **C)** Top 20 upstream regulators of DEGs in all B cells of SjD patients and non-SjD controls using the IPA regulator database. **D-F)** UMAP plots colored by expression of the GEPs associated with B cell identify **(D)**, naive B cells **(E)**, and SjD specific genes in B cells **(F)** identified by NMF. The bar plots show usage scores for these GEP in naive, intermediate and memory B cells of SjD patients and non-SjD controls. **G)** Top 10 MGI (mouse genome informatics) phenotypes associated with the SjD B cell GEP identified by EnrichR analysis. **H)** Heatmap of DEGs contained in the *SjD B cell* GEP in all B cells of SjD patients and non-SjD controls. Colors depict log_2_ fold changes (FC) in expression. Statistical analysis in (A), (D), (E), and (F) was performed using a Wilcoxon rank sum test. The significance of pathways and regulators in (B) and (C) was calculated using a right-tailed Fisher’s exact test. The significance in (A), (B), (C), (D), (E), and (F) was adjusted using the Benjamini-Hochberg method. Pathways shown in (G) are the top 10 ranked by P value identified using a right-tailed Fisher’s exact test. Scales shown in (D-F) represent the usage scores per cell for each GEP with a range between 0.2 (low) and 0.7 (high). *P < 0.05; **P < 0.01; ***P < 0.001.

To identify functional gene expression profiles that correlate with specific B cell subsets and are used specifically by patients with SjD, we again conducted an NMF analysis of all B cell subsets of patients and controls. We identified three GEPs, of which one was ubiquitously used by all three B cell subsets, and we therefore called *B cell identity* GEP **(Figure 3D)**. It contains the canonical B cell markers we used to identify B cells by scRNA-seq (**Figure 1A**). Naive B cells of SjD patients showed a significant usage of this GEP compared to non-SjD controls, although the differences were modest. A second GEP we identified in B cells was predominantly used by naïve B cells of both patients and controls, whereas intermediate and memory B cells do not utilize this GEP (**Figure 3E**). Usage of this *Naive B cell* GEP was significantly depleted in naïve B cells from SjD patients compared to non-SjD controls. A third GEP was found in all three B cell subsets and different from the other two B cell GEPs because its usage was significantly increased in naive, intermediate, and memory B cells of SjD patients compared to controls (**Figure 3F**). The overall usage of this *SjD B cell* GEP was highest in naive B cells. Because naive B cells were also the only subsets whose numbers were significantly increased in SjD patients **(Figure 3A)**, we hypothesized that the genes contained in this GEP might be responsible for the increased B cell frequencies in patients.

A functional enrichment test of the genes contained in the *SjD B cell* GEP using EnrichR identified genes that are associated with phenotypes related to B cell physiology, proliferation, and morphology, impaired humoral immune responses, and decreased transitional B cell numbers **(Figure 3G)**. Surprisingly, we also identified genes associated with increased T cell numbers, especially memory CD8^+^ T cells, although we had not observed an increased abundance of CD8^+^ T cells in SjD patients **(Figure 1A)**. We analyzed the differential expression of genes belonging to the *SjD B cell* GEP in naive B cells from SjD patients and controls **(Figure 3H**). The vast majority of DEGs are enriched in B cells of SjD patients with the most highly enriched genes including *ROR1* (encoding Receptor Tyrosine Kinase-Like Orphan Receptor 1), which plays a role in the development of B cells, and *GAB1* (GRB2-Associated Binder 1), a protein that regulates proliferation via PI3K/AKT and MAPK/ERK pathways, although its role in B cells is not known. Enriched genes also include known regulators of B cell fate and function, including *MZB1* (encoding marginal zone B and B1 cell-specific protein), *SYK* and *LYN* (encoding kinases required for B cell receptor signaling), *PAX5* and *BACH2* (transcription factors regulating different stages of B cell development). Taken together, we find an expansion of naive B cells in patients with SjD and identify increased usage of a GEP in naive B cells of patients that contains many genes involved in B cell development and function. These data provide indirect evidence that naive B cells from SjD patients are activated in a disease-specific manner.

### Depletion of a signal transduction GEP in CD8^+^ T cells of SjD patients

CD8^+^ T cells contribute to injury of acinar or ductal epithelial cells in the exocrine glands of patients SjD by a variety of mechanisms, including the production of IFN-γ and induction of apoptosis ^50^. A recent study showed that granzyme K-producing CD8^+^ T cells target seromucous acinar cells in patients with SjD ^51^. CD8^+^ T cells showed the strongest differential gene expression with more than 4,500 genes up- or downregulated in SjD patients compared to controls **(Figure 1D)**, suggesting that this T cell subset is significantly functionally dysregulated in patients’ PBMC. Moreover, CD8^+^ T cells from patients showed a profound depletion of translation and ribosome-associated pathways without associated changes in the frequencies in CD8^+^ T cells within PBMCs of SjD patients **(Figure 1C,E)**. To understand the functional state of CD8^+^ T cells, we first conducted a pathway analysis of DEGs in SjD patients compared to controls using the IPA platform. We observed a strong enrichment of type I and type II IFN signaling pathways, as well as TCR signaling, Th1 pathway, and death receptor signaling in the patients’ CD8^+^ T cells **(Figure 4A)**. Of note, while depletion of ribosome-associated pathways was again observed, it did not appear as one of the top dysregulated pathways. Consistent with the pathway analysis, we identified many interferons and IFN signaling-related factors as drivers of gene expression in CD8^+^ T cells of SjD patients using an upstream regulator analysis **(Figure 4B)**. These included IFN-α, IFN-γ, IFNL1, IRF7 and STAT1. These findings related to enhanced IFN signaling are similar to those in other immune cell subsets **(Figure 1E)** and likely are secondary to the overall IFN-driven inflammatory milieu in SjD patients. To identify additional changes in gene expression that are more specific to CD8^+^ T cells and that may provide insight into their functional state, we again used NMF. We identified a GEP that is specific to a subpopulation that corresponds to naive CD8^+^ T cells and is not used by CD4^+^ T cells or other T cells either **(Figure 4C)**. We therefore named it *Naive CD8^+^ T cell* GEP. The usage of this GEP was significantly depleted in CD8^+^ T cells of patients with SjD. A functional enrichment analysis of the genes in this GEP revealed that many of them are associated with TCR signaling (**Figure 4D**). Specific pathways these genes were linked to included beta-catenin, canonical NF-kB, calcineurin-NFAT, and Ras signaling. As expected from the reduced usage of this GEP, the expression of almost all genes was reduced in CD8^+^ T cells of SjD patients **(Figure 4E)**. Some of the genes in this GEP whose expression was most strongly depleted and that have been associated with CD8^+^ T cell function before were *S100B, CD37, CCR7,* and *RCAN3*. Many of the depleted genes have either positive or negative regulatory roles in TCR signaling and CD8^+^ T cell function, and it is, therefore, difficult to determine the functional consequences resulting from the reduced usage of this GEP by CD8^+^ T cells of SjD patients.

**Figure 4.**
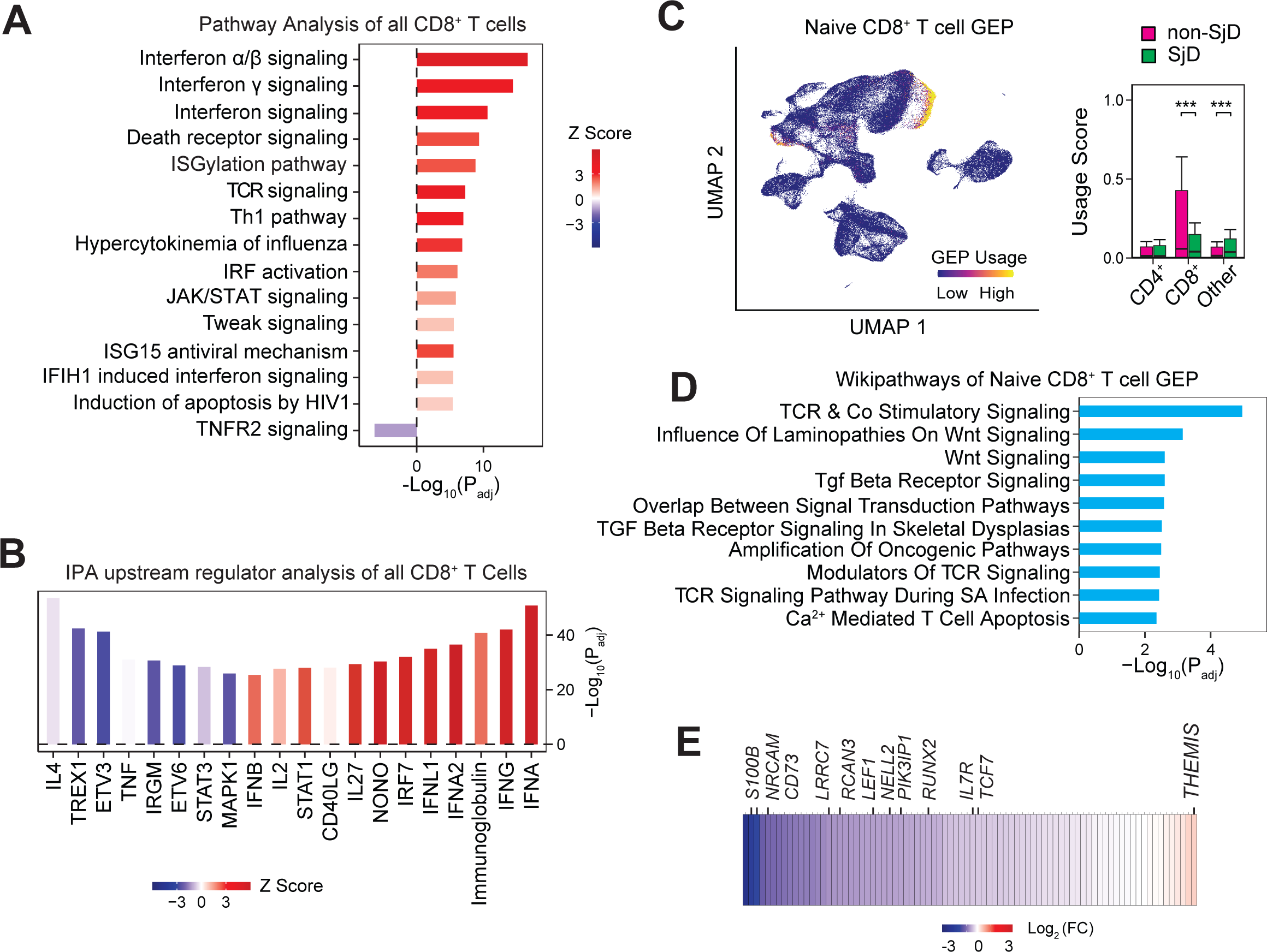
Depletion of genes in a GEP associated with naive CD8^+^ T cells in patients with SjD. **A)** Top 15 dysregulated pathways based on DEGs in all CD8^+^ T cells of SjD patients and non-SjD controls using the canonical pathways of the IPA platform. **B)** Top 20 upstream regulators of DEGs in all CD8^+^ T cells of SjD patients and non-SjD controls using the IPA regulator database. **C)** UMAP plot of PBMCs colored by expression of the GEP associated with naive CD8^+^ T cells identified by NMF. The bar plot shows usage scores for this GEP in all CD4^+^ T cells, CD8^+^ T cells and other T cells of SjD patients and non-SjD controls. **D)** Top 10 NCI Nature pathways associated with the *naive CD8^+^ T cell* GEP identified by EnrichR analysis. **E)** Heatmap of DEGs contained in the *naive CD8^+^ T cell* GEP in all CD8^+^ T cells of SjD patients and non-SjD controls. Colors depict log_2_ fold changes (FC) in expression. Statistical analysis in (C), and (D) was completed using a Wilcoxon rank sum test. The significance of pathways and regulators shown in (A) and (B) was calculated using a right-tailed Fisher’s exact test. The significance in (A), (B), (C), and (D) was adjusted using the Benjamini-Hochberg method. The scale in (C) represents the usage scores per cell for the GEP with a range between 0.2 (low) and 0.7 (high). *P < 0.05; **P < 0.01; ***P < 0.001.

### Two distinct functional GEPs in CD4^+^ T cells of SjD patients and signs of enhanced Th1 function

CD4^+^ T cells have been extensively studied in SjD, and they make up a large portion of the lymphocytic infiltrates found in SGs of patients ^52^. Several different CD4^+^ T cell subsets, including T helper (Th) 1, Th17, and T follicular helper (Tfh) cells, have been implicated in SjD pathogenesis ^52^. Another CD4^+^ T cell subset relevant to SjD is regulatory T cells (Treg), whose frequencies in the blood and exocrine glands of SjD patients have alternatively been reported to be increased, decreased, or unchanged ^53–62^. We recently reported enhanced functional gene expression signatures in Th1 cells and decreased signatures in Treg cells in the blood of patients with SjD ^63^. Here, we assessed differential gene expression in all CD4^+^ T cells of patients with SjD and non-SjD controls to identify potential imbalances in the different subsets of CD4^+^ T cells and gain insights into their functional state. A pathway analysis of DEGs in all CD4^+^ T cells revealed the enrichment of pathways related to TCR signaling, oxidative phosphorylation (OXPHOS), and the regulation of apoptosis **(Figure 5A)**. These findings were confirmed by GSEA (**Figure 5B**). Enriched genes in CD4^+^ T cells of SjD patients included well-known molecules involved in TCR signaling, including *CD3E, CD3G, LCK, ZAP70*, and *LAT* **(Figure S4B)**. Although not detected by IPA pathway analysis, our manual analysis and GSEA of enriched genes also identified many *PSM* genes that encode proteasome subunits, which may indicate increased protein degradation in CD4^+^ T cells of SjD patients **(Figure S4B, C)**. By contrast, pathways related to protein translation were strongly downregulated in patients’ CD4^+^ T cells **(Figure 5A)**. An upstream regulator analysis of DEGs using the IPA platform identified many T cell-associated surface receptors and cytokines to be enriched in SjD patients **(Figure 5C)**. These included the TCR, CD3, CD28, and CD40L, as well as IL-2, IFN-γ and IL-15. IFN-γ is the signature cytokine of Th1 cells, and IL-15 promotes the proliferation of developing Th1 cells (especially in combination with IL-12), potentially indicating enhanced polarization of CD4^+^ T cells into Th1 cells in SjD patients. The upstream regulator analysis also found a strong enrichment of LARP1 and depletion of MYC in CD4^+^ T cells of SjD patients, which is similar to findings in their B cells and correlates with the depletion of genes and pathways associated with protein translation.

**Figure 5.**
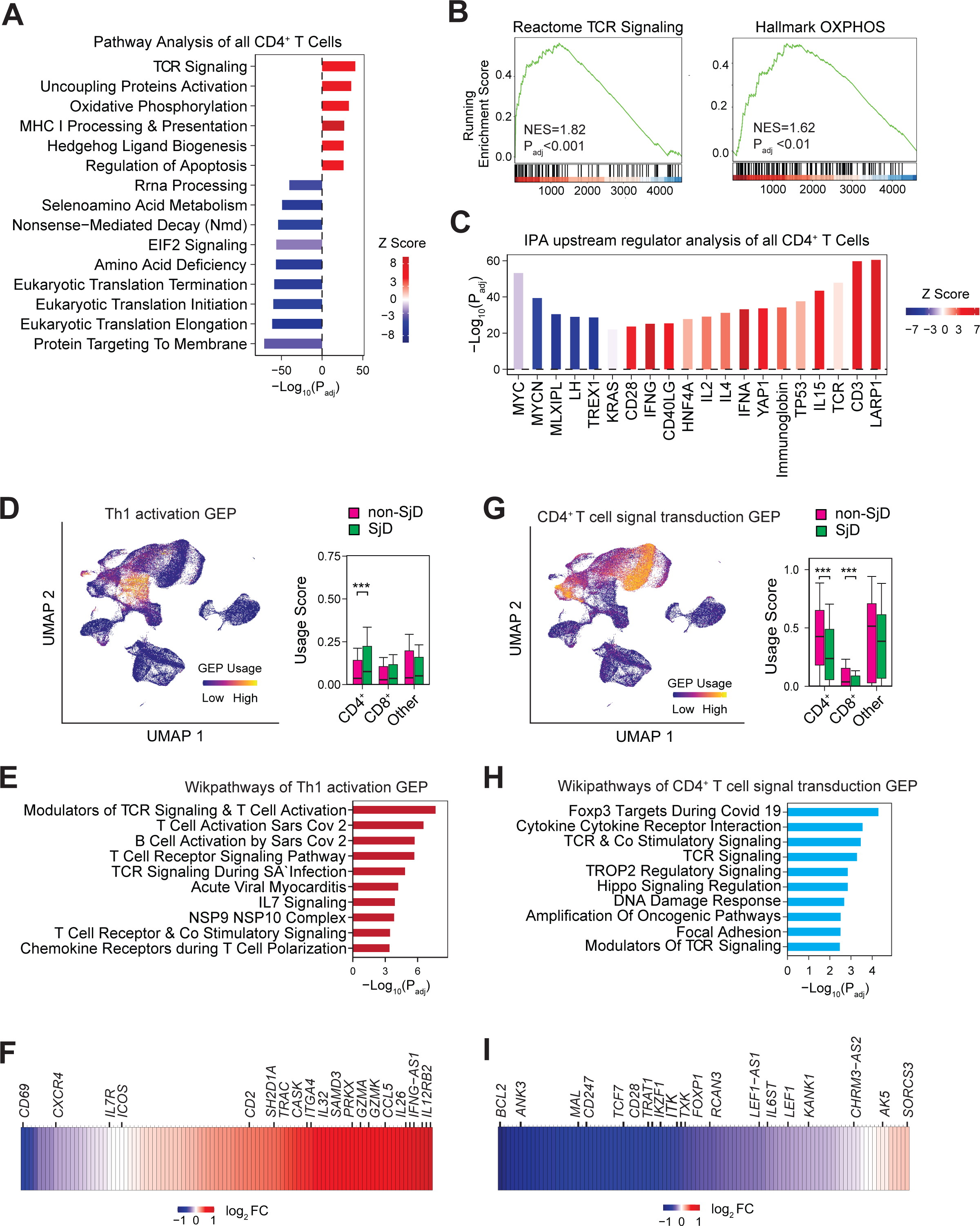
Enhanced usage of a functional GEP in CD4^+^ Th1 cells of SjD patients. **A)** Top 15 dysregulated pathways based on DEGs in all CD4^+^ T cells of SjD patients and non-SjD controls using the canonical pathways of the IPA platform. **B)** GSEA of DEGs in all CD4^+^ T cells of SjD patients and controls. Shown are enriched gene sets related to TCR signaling and OXPHOS. **C)** Top 20 upstream regulators of DEGs in all CD4^+^ T cells of SjD patients and non-SjD controls using the IPA regulator database. **D)** UMAP plot of PBMCs colored by expression of the GEP associated with Th1 activation identified by NMF. The bar plot shows usage scores for this GEP in all CD4^+^ T cells, CD8^+^ T cells and other T cells of SjD patients and non-SjD controls. **E)** Top 10 Wikipathways associated with the *Th1 activation* GEP identified by EnrichR analysis. **F)** Heatmap of DEGs contained in the Th1 activation GEP of all CD4^+^ T cells of SjD patients and non-SjD controls. Colors depict log_2_ fold changes (FC) in expression. **G)** UMAP plot of PBMCs expressing a GEP associated with CD4^+^ T cell signal transduction identified by NMF. The bar plot shows usage scores for this GEP in all CD4^+^ T cells, CD8^+^ T cells and other T cells of SjD patients and non-SjD controls. **H)** Top 10 Wikipathways associated with the *CD4^+^ T cell signal transduction* GEP identified by EnrichR analysis. **I)** Tile plot of DEGs contained in the CD4^+^ signal transduction GEP of all CD4^+^ T cells of SjD patients and non-SjD controls. Colors depict log_2_ fold changes (FC) in expression. Statistical analysis in (D), (E), (G), and (H) were performed using a Wilcoxon rank sum test. The significance of pathways and regulators shown in (A) and (C) was calculated using a right-tailed Fisher’s exact test. The significance in (A), (B), (C), (E), (G), and (H) was adjusted using the Benjamini-Hochberg method. Scales shown in (D) and (G) represent the usage scores per cell for each GEP with a range between 0.2 (low) and 0.7 (high). *P < 0.05; **P < 0.01; ***P < 0.001.

We next conducted an NMF analysis of all CD4^+^ T cells to identify functional gene expression profiles that correlate with specific CD4^+^ T cell subsets and may be used specifically by SJD patients. We identified two major GEPs in CD4^+^ T cell subsets **(Figure 5D, G)** as identified by the expression of marker genes **(Figure S1A)**. One was predominantly utilized by CD4^+^ T cells that were identified as Th1 cells by marker gene expression **(Figure S1A)**. Its usage was significantly increased in CD4^+^ T cells (but not other T cell types) of SjD patients compared to controls **(Figure 5D).** To identify the biological function of this GEP, we performed a functional enrichment test on the genes contained within it (**Figure 5E**). Most of the significantly enriched pathways associated with genes in this GEP were linked to TCR signaling, suggesting that this GEP is related to T cell activation. Individual inspection of DEGs contained in this GEP revealed that most were enriched in SjD patients. These included *IL-12RB2* (a subunit of the IL-12 receptor that is required for Th1 differentiation) and *IFNG-AS1* (a long noncoding RNA, lncRNA, also called TMEVPG1, that is associated with increased Th1 responses in SjD ^64^) (**Figure 5F, Figure S4D**). We thus named this GEP *Th1 activation*. Our findings suggest that Th1 cells are hyperactivated in the blood of patients with SjD.

The other GEP we identified was expressed in different CD4^+^ T cell subsets **(Figure 5G, Figure S1A)**. Although broadly expressed in CD4^+^ T cells, this GEP does not overlap with the *Th1 activation* GEP, indicating that they are functionally distinct from one another. The usage of the second GEP was significantly depleted in CD4^+^ T cells of SjD patients compared to non-SjD controls **(Figure 5G)**. Upon performing a FET to determine the biological implications of these genes being depleted, we identified pathways related to signal transduction, generation of second messengers, WNT signaling, and TCF/LET target gene regulation to be linked with this GEP **(Figure 5H)**. Individual inspection of genes in this GEP revealed that most of them were depleted in CD4^+^ T cells of SjD patients **(Figure 5I, Figure S4E)**. These genes include regulators of signal transduction in T cells such as *CD247* (encoding the TCRζ chain), *ITK, CAMK4,* and *CD28*, regulators of T cell development including *THEMIS*, *TCF7,* and *LEF1* and genes promoting T cell survival such as *BCL2* and *IL7R*. The functional significance of gene depletion in this *CD4^+^ T cell signal transduction* GEP is unknown.

## DISCUSSION

Compared to its prevalence, the pathophysiology of SjD is inadequately understood. High throughput techniques such as scRNA-seq allow for unbiased, granular analyses of the prevalence of different immune cell types in SjD patients and their functional status. Here we show that scRNA-seq analysis, in conjunction with machine learning based approaches, is able to identify functional gene expression profiles (GEPs) in PBMCs of patients with SjD that may correlate with the function of immune cells during disease.

Cell type identification by scRNA-seq in PBMCs of patients with severe SjD (ACR/EULAR scores of 5-9) compared to non-SjD patients revealed an increase in the frequency of B cells and a decrease in the frequency of CD4^+^ T cells, whereas those of other immune cells were unchanged. The increase in B cells was driven mostly by naive B cells, not memory B cells whose frequency was unchanged. B cells have long been implicated in the pathophysiology of SjD as emphasized by the presence of autoantibodies and the fact that B cell infiltrates are found in SGs of patients with SjD. B cells were shown to be hyperactive in SjD, resulting in increased proliferation and antibody production^65^. Here we identify several functional GEPs in B cells whose usage is altered in patients with SjD. A GEP corresponding to naive B cells is depleted in SjD patients, whereas a GEP that is specifically associated with B cells in SjD patients is increased compared to non-SjD controls. The latter *SjD B cell* GEP contained genes involved in both B cell activation and proliferation. Most of the genes in this B cell signature were upregulated, including the tyrosine kinases Lyn and Syk which regulate the proliferation and activation of B cells^66,67^. In SjD, treatment with remibrutinib, a Bruton’s tyrosine kinase (BTK) inhibitor, has been shown to alleviate symptoms, and reduce serum levels of SSA/B in SjD patients^68^. These findings underscore the important role of B cells in SjD and the effectiveness of monoclonal antibodies that deplete B cells (anti-CD20, rituximab) or inhibit B-cell activating factor (BAFF) signaling (belimumab, ianalumab) in alleviating SjD symptoms ^69,70^. Our findings are consistent with previous reports of B cell hyperactivation in SjD and show that they utilize a unique GEP whose constituent genes may be new therapeutic targets in SjD.

CD4^+^ T cells play an important role in the pathophysiology of SjD and their numbers are found to be increased in the salivary and lacrimal glands of SjD patients^52^. CD4^+^ T cells can be divided into distinct T helper (Th) cell subsets, including Th1, Th17 and T follicular helper (Tfh) cells, all of which have been implicated in SjD^71^. Tfh cells help B cell to differentiate into antibody producing plasma cells or memory B cells. Increased frequencies of Tfh cells were found in the blood and glands of patients with SjD^72–75^ and correlated positively with ESSDAI scores^76,77^. Th17 cells are involved in several autoimmune diseases and while most studies did not find aberrant frequencies of IL-17^+^ cells in the blood of patients^74,78,79^, IL-17 mRNA levels were increased in their MSG^80,81^. Th1 cells, whose major effector cytokine is IFN-γ, play important pathogenic roles in several autoimmune diseases^82,83^ including SjD^84–87^. Here we report that the frequency of CD4^+^ T cells among PBMC of SjD patients is decreased. CD4^+^ T lymphopenia in the blood of patients is a common laboratory finding in SjD. Reduced CD4^+^ T cell counts are correlated with higher systemic disease activity scores and are considered a predictive factor for lymphoma development ^38,39,88^. The reduction of CD4^+^ T cells we observed was not driven by the depletion of a single subtype of CD4^+^ T cells of the twelve we identified in PBMCs by scRNA-seq. As we reported recently, the frequencies of 11 CD4^+^ T cell subsets were comparable between SjD patients and non-SjD controls including Th1, Th17 and Tfh cells^63^. The only exception was type 1 regulatory (Tr1) cells whose frequencies were increased. Because the frequencies of most CD4^+^ T cell subsets were normal in SjD patients, we investigated their functional gene expression signatures. NMF identified two distinct GEPs in CD4^+^ T cells of SjD patients. One of these GEPs, which we named *Th1 activation* GEP, was associated mostly with Th1 cells and its utilization was increased in CD4^+^ T cells of SjD patients. Most genes in this GEP showed increased expression in SjD patients including several that had previously been linked to Th1 function, including *IL-12RB2* (a subunit of the IL-12 receptor required for Th1 differentiation), *IFNG-AS1* (a lncRNA associated with increased Th1 responses in SjD ^64^) and the proinflammatory cytokine *IL26.* Our finding of hyperactivated Th1 cells is consistent with a recent study showing that ∼ 30% of CD4^+^ T cells in salivary glands of SjD patients express the transcription factor T-bet, identifying them as Th1 cells^89^. Moreover, the expression of IFN-γ is increased in SGs and saliva of patients with SjD^84–87,89–91^, and IFN-γ expression correlated with the degree of lymphocytic infiltration^86^.

This hyperactivation was specific to Th1 cells and not a general feature of all CD4^+^ T cells. A second GEP we identified in CD4^+^ T cells, named the *CD4^+^ T cell signal transduction* GEP, was significantly depleted in the vast majority of SjD patients’ CD4^+^ T cells except Th1 cells. Genes in this GEP are associated with pathways linked to signal transduction and most of these genes were depleted including *CD247* (encoding the TCRζ chain), *ITK, CAMK4* and *CD28*. Moreover, we found genes related to T cell development (such as *THEMIS*, *TCF7* and *LEF1*) and T cell survival (such as *BCL2* and *IL7R*) to be depleted. We made a similar observation in CD8^+^ T cells of SjD patients. The only GEP we identified that is specifically used by CD8^+^ T cells, or more precisely naive CD8^+^ T cells, was significantly depleted in SjD patients. This *naive CD8^+^ T cell* GEP contains many genes that are linked to TCR signaling including NF-kB, calcineurin-NFAT and Ras signaling. The expression of almost all genes within this GEP was reduced in CD8^+^ T cells of SjD patients including *S100B, CD37, CCR7* and *RCAN3*. Some of the depleted genes overlap with those found in the *CD4^+^ T cell signal transduction* GEP including *TCF7, LEF1 and RCAN3*. Because the depleted genes in both CD4^+^ and CD8^+^ T cell GEPs have either positive or negative regulatory roles in TCR signaling and T cell function, the functional significance of their depletion remains to be determined.

Multiple recent studies have used scRNA-seq to investigate cell populations and their functional states in PBMCs and salivary glands of patients with SjD. Several of these studies have identified CD8^+^ cytotoxic T cells (CD8^+^ CTLs) infiltrating the SGs of patients with SjD ^26,30,32,51,89^, which were reported to express the serine protease GZMK^30,32,51^ or CD9^26^ endowing them with cytotoxic and tissue-resident properties, respectively. In the peripheral blood, GZMK^+^CXCR6^+^CD8^+^ T cells were increased and harbored a gene signature similar to tissue-resident memory T cells in SGs of SjD patients^32^. Intriguingly, CD8^+^ T cells in the blood and tissue-resident labial SGs of patients shared similar TCR clones^32^. Together, these reports provide good evidence for the presence of tissue-resident cytotoxic CD8^+^ T cells in SGs of SjD patients. Unlike the study by Xu et al.^32^ we did not observe increased cytotoxic CD8^+^ T cells numbers in the blood of patients with SjD. We did, however, identify a unique GEP that was associated with naive CD8^+^ T cells whose utilization was reduced in SjD patients. Many of the depleted genes in this *naive CD8^+^ T cell* GEP regulate TCR signaling and function, indicating that circulating naive CD8^+^ T cells in the blood of SjD patients may be dysfunctional.

Besides CD8^+^ CTL, one study showed an increase in the proportion of CD4^+^ T cells in the peripheral blood of SjD patients that express GZMB marking them as potentially cytotoxic ^24^. We also detected GZMB-expressing CD4^+^ CTL by scRNA-seq but their numbers were comparable in SjD patients and controls ^63^. Of note, the study by Hong et al.^24^ did not identify several other CD4^+^ T cell subtypes including Th1 and Treg cells that have been implicated in SjD pathology and are regularly detected in PBMC by scRNA-seq^92^. In fact, none of the other scRNA-seq studies have identified a Th1 signature in CD4^+^ T cells of patients with SjD, either in the blood or in SGs. Neither did we detect Th1 cells in our analysis of PBMCs from SjD pathways using conventional pathway or upstream regulator analyses of DEG. We identified a GEP associated with Th1 cells in SjD patients by using NMF. Usage of this *Th1 activation* GEP was increased in SjD patients driven by the enhanced expression of the majority of genes in this GEP. Besides Th1 associated genes, this GEP contained *GZMB* and *GZMK* whose expression was increased. It is possible that this GEP has some functional overlap with CD4^+^ CTL. As discussed above, our identification of a *Th1 activation* GEP in SjD patients is validated by a recent study reporting that ∼ 30% of CD4^+^ T cells in SGs of SjD patients are Th1 cells^89^ and the fact that IFN-γ levels are increased in SjD patients^84–87,89–91^. In most scRNA-seq studies discussed above altered B cell numbers or phenotypes are not a prominent finding. Xiang et al.^31^ report expanded IGHD^+^ naive B cells in the blood of SjD patients and increased numbers of memory, effector memory and naive B cells in their SGs. Our study also shows an increase in B cell numbers in the blood of SjD patients that is mainly driven by an expansion of naive IGHM^+^IGHD^+^ B cells. However, naive B cells in SjD patients show reduced usage of a GEP that is specifically associated with naive B cells. Instead, they appear to utilize a GEP that is specific to SjD patients suggesting that naive B cells in SjD patients are activated in a disease specific manner.

Like several of the other scRNA-seq studies, we observed a strong increase in the expression of IFN induced genes and IFN signaling pathways that was apparent in all immune cell populations we analyzed^24,27,28,30^. These genes included IFI44L, ISG15, STAT1 and others. While this increase is likely secondary to elevated levels of type I and type II interferons in SjD patients, we were not able to identify a single cell type as the source of IFN production in the blood of patients including pDCs and monocytes. A unique finding of our study is the strong depletion of genes encoding ribosomal proteins and pathways related to translational initiation and elongation. These changes, which were specific to CD4^+^ and CD8^+^ T cells as well as B cells, have to our knowledge not been reported in SjD before. Besides a depletion of pathways related to protein synthesis, we also observed an increased expression of *PSM* genes encoding proteasome subunits in CD4^+^ T cells and monocytes, potentially resulting in enhanced protein degradation in SjD patients. Collectively these findings suggest that protein homeostasis is perturbed in lymphocytes of SjD patients.

A potential limitation of our study is the fact that we analyzed PBMCs and not immune cells infiltrating the SGs of SjD patients, which might provide a more direct picture of the functional state of these cells. To compare immune cells in inflamed SGs and blood of SjD patients, researchers have used paired samples of both glandular tissue and PBMCs with to goal to assess the functional state of gland-infiltrating immune cells without the need for tissue resection. These studies have shown that changes in gene expression, in particular the IFN-α signture, in immune cells in the blood and SGs correlate well^93^, validating the utility of immune cell characterization in the peripheral blood of SjD patients.

Although several studies have analyzed PBMC and SGs of SjD patients by scRNA-seq, our study is the first to our knowledge to utilize machine learning approaches to analyze GEPs in SjD patients’ cells. NMF has allowed us to identify unique GEPs that were not detected by standard pathway analyses. These findings include the identification of a unique *Th1 activation* GEP that is utilized more strongly in SjD patients than symptomatic non-SjD controls. We also detected two GEPs in CD4^+^ and CD8^+^ T cells that contain genes involved in T cell activation, which are depleted in SjD patients. Moreover, we identified a GEP that was specifically utilized by B cells of SjD patients, particularly naive B cells. This GEP may be involved in the abnormal activation and proliferation of B cells, a phenomenon commonly observed in SjD. These findings demonstrate that NMF is able to provide new insights into the pathophysiology of SjD and, by extension, other autoimmune disorders. Moreover, these results may help identify novel drug targets for the treatment of SjD.

## MATERIALS AND METHODS

### Patient samples

Patient data and peripheral blood mononuclear cell (PBMC) samples of patients with SjD and non-SjD controls were obtained from the Sjögren’s International Collaborative Clinical Alliance (SICCA)^94,95^. The age, gender, autoantibody status, focus score and the 2016 ACR-EULAR SjD classification criteria score are shown in **Table S1**. The collection of patient samples by SICCA was approved by the University of California, San Francisco IRB committee (IRB 10-02551). Participants gave informed consent to participate in the study before taking part in the study.

### Single-cell RNA sequencing of PBMC

PBMCs were thawed and stained with Annexin-V (Biolegend, Cat. 640912) and LIVE/DEAD (ThermoFisher, Cat. L23105) in Annexin-V binding buffer (Biolegend, Cat. 422201). Live PBMC were sorted as Annexin-V^−^ Live/Dead^−^ on an Aria II (BD Biosciences) with a 70 μm nozzle. Live cells were collected in RPMI1640 medium (Corning, Cat. 10040CV) with 10% fetal bovine serum. Sorted cells were stained using TotalSEQ Type B hashing antibodies (BioLegend, Cat. 394633 to 394649) for patient identification post multiplexing (**Table S2**). Collected cells were quantified using a Bio-Rad TC20 automated cell counter. Hash-Tag antibody labeled cells were encapsulated into emulsion droplets using the 10x Genomics platform for library preparation. scRNA-seq and antibody multiplexing libraries were constructed using the Chromium Single Cell 3’ v3.1 Reagent Kit and 3’ Feature Barcode Kit (PN-1000268 and PN-1000262) according to the manufacturer’s protocols. Individual libraries were diluted to 2 nM and pooled for sequencing. Pooled batches were sequenced with 100 cycle run kits (28 bp Read1, 8 bp Index1 and 91 bp Read2) on the NovaSeq6000 Sequencing System (Illumina). Amplified cDNA was evaluated with an Agilent BioAnalyzer 2100 using a High Sensitivity DNA Kit (Agilent Technologies). Final libraries were evaluated with an Agilent TapeStation 4200 using High Sensitivity D1000 ScreenTape (Agilent Technologies).

### Single-cell transcriptome assembly

For each batch of patients, reads from the Illumnia platform were converted to fastq files using the bcl2fastq package. Fastq files were processed into transcriptome assemblies using 10x Genomics Cell Ranger 7.0.0 and assembled using the GRCh38 Genome^96^. Hashtag values were assigned to each cell in tandem to assembly, and demultiplexing was completed for each batch prior to integration. All analyses were performed using R Statistical Software (v4.1.2; R Core Team 2021) unless otherwise noted. Downstream single cell analysis was performed using Seurat (v5.0.1)^97^. Visualization and figure generation was performed using ggplot2 (v3.4.4) unless otherwise noted^98^.

### Demultiplexing and quality control of cells

Prior to combining the three batches of scRNA-seq data into one Seurat object, each batch was subjected to quality control and demultiplexing. Hashtag counts were normalized using a centered log-ratio method, and positive cells were defined via Seurat’s HTOdemux function, using the 0.99 quantile as the positive cut-off. Cells identified as doublets were filtered out from further downstream analysis. After doublet removal, cells were further filtered based on mitochondrial content and feature content. Cells with greater than 15% of features being mitochondrial genes were removed; likewise cells with less than 1,000 unique molecular identifiers (UMIs) were removed. Cells falling into the 98^th^ and 2^nd^ percentiles for the number of genes covered per cell were also filtered as doublets and negatives, respectively. After quality control for doublet, singlet, and mitochondrial percentage, the remaining cells were integrated using Seurat with good mixing of data, and a robust number of variable genes. A total of 105,152 cells were successfully demultiplexed, and subsequently passed quality control filters, with a mean of 6,185 cells per sample with a similar distribution across all batches.

### Normalization and integration of samples

Gene expression levels were normalized per cell by log normalization and then scaled by a factor of 10,000. To correct for potential batch effects and sampling effects, each patient was individually integrated using variance stabilizing transformation (SCTv2), setting both batch and mitochondrial percentages as variables to regress out. After transformation, samples were integrated by reciprocal principal component analysis (RPCA) using 4,000 highly variable features^99^. To address potential disease state bias, one non-SjD control sample and one SjD patient sample were selected as integration anchor references. Both samples were from the same batch, and they were selected for having the highest average UMI per cell.

### Cell clustering and cell type annotation

To identify different immune cell types, we used the integrated assay of all combined samples and performed a PCA calculating the first 75 components. We then generated a uniform manifold approximation projection (UMAP) and a t-distributed stochastic neighbor embedding (t-SNE) plot using the first 60 dimensions and 20 nearest neighbors. Cell clusters were calculated with the same settings as both dimensional reduction plots, with a resolution of 1.4. Calculated clusters were annotated using SingleR, using the Database Immune Cell Expression Data as a reference^36,100^. The azimuth reference was used as a second database with similar cell type identification^97^. Separate clusters that matched to the same cell type were combined into a singular cluster. These combined clusters were then confirmed using known marker genes for each cell.

### Analysis of cell type annotation using cell type markers

To confirm that we had properly identified each cluster and their corresponding cell types, we grouped the cells on two different levels of granularity. The first, broader-level annotation grouped cells into seven broad cell types, defined as CD4^+^ and CD8^+^ T cells, other T cells that were neither CD4^+^ nor CD8^+^, natural killer (NK) cells, B cells, monocytes, and dendritic cells (DCs). We used canonical cell type markers listed in the Azimuth database for cell type clustering as follows: Monocytes (CTSS, FCN1, NEAT1, LYZ, PSAP, S100A9), DCs (CD74, HLA-DPA1, HLA-DPB1, HLA-DQA1, CCDC88A, HLA-DRA, HLA-DMA), B cells (CD79A, RALGPS2, CD79B, MS4A1, BANK1, CD74), NK cells (GNLY, GZMB, NKG7, KLRF1, PRF1), CD4+ T cells (IL7R, MAL, LTB, CD4, LDHB), CD8+ T cells (CD8A, CD3D, TMSB10, HCST, CD3G, LINC02446), and other T cells (TRDC, GZMK, IL7R, ANK3). The cellular composition of each sample was calculated using the total number of filtered cells per sample, and the mean cellular frequency was calculated for each disease state. Within each of these seven broad annotations, more granularly resolved cell types were confirmed using the appropriate marker genes, as illustrated in the corresponding supplemental figures **(Figure S2A,B** and **Figure S3A,B)**. Marker genes for both broad annotations and granular subtypes are examples and are non-exhaustive; additional canonical markers were used to confirm proper cluster identification. Results are included in Supplemental Data File 1A.

### Cellular communication analysis

To determine if cell-to-cell communication between the seven major immune cell subsets within PBMCs was perturbed in SjD patients, we used the Cellchat package(v1.6.1)^30^. We accessed the CellchatDB for the collection of ligand-receptor pairs using the secreted signaling and ECM Receptor sections. Cell-to-cell communication was calculated independently for SjD and non-SjD control cohorts using the same pipeline, database, and default thresholds. The false discovery threshold for cellular interactions for each disease state was set at a Bonferroni-adjusted value (Padj) ≤ 0.01. The bottom 50^th^ percentile of weighted discovered ligand-receptor pairs was removed due to low gene expression levels. After calculating cell-to-cell communication for each condition, we subtracted the normalized values for each ligand-receptor pair to determine differential communication.

### Non-negative matrix (NMF) factorization

To identify unique cell-type functions and their changes between SjD patients and non-SjD controls, we performed consensus non-negative matrix factorization (cNMF) as described in^45^. cNMF is based on the NMF implementation in scikit-learn v0.20.0. We filtered out genes that were not expressed in at least 20 cells across all samples and removed any cells that had less than 500 genes. No cells were lost via this filtering. We then calculated the top 3,000 overdispersed genes and used only those to perform NMF analysis. To determine the number of components (k) that would be most accurate, we calculated k over a range of 10 to 50, with each value having 200 iterations. We calculated the silhouette score and reconstruction error as implemented in cNMF. We found that k = 34 is the smallest, most stable solution. The final consensus solution was determined using a density threshold of 0.1 to exclude outlier solutions. We identified a total of 20 gene expression profile (GEP) usages with a mean score of greater than 0.1 for any cell type. The remaining 14 usages (of 34 total) were not commonly used by any cell type or groups of cell types. Marker genes for each GEP were calculated using multiple least squares regression of normalized z-scored gene expression against the consensus GEP usage matrix as implemented by cNMF. A functional enrichment test for differentially utilized GEP was subsequently performed using the top 100 marker genes per GEP using the EnrichR platform^101^. cNMF analysis was performed in Python (v.3.7.0) using scanpy (v.1.6.0), pandas (v.1.1.3), numpy (v.1.19.2), matplotlib (v.3.3.2)^102,103^. After cNMF analysis, the scanpy object with corresponding per cell values for each GEP, was converted back into Seurat, for subsequent analysis and quantification.

### Differential gene expression analysis

To identify differentially expressed genes (DEGs) between SjD patients and non-SjD donors, we divided each identified cell type cluster by disease state. All cell types were confirmed to have equal representation in batches. We then calculated differential gene expression between the disease states using the FindMarkers function in Seurat. The RNA assay was utilized for expression analysis as per Seurat publisher’s recommendation^92^. When calculating DEGs we used the MAST algorithm implemented by Seurat^41^. We considered all genes found in at least 10% of cells with an absolute log_2_ fold-change (FC) ≥ 0.1 and Padj ≤ 0.01 to be significant. Non-significant expression changes were calculated as well to act as background genes for downstream gene set enrichment analysis (GSEA).

### Functional enrichment analysis

To determine the functional enrichment or depletion of pathways associated with DEGs in each cell type, we performed pathway analyses. Using DEGs between SjD patients and controls, we performed a functional enrichment test (FET) using the GSEA function of ClusterprofileR 4.0^42^. Databases queried for our GSEA analysis included a combination of the canonical databases of molecular signature database, along with all 3 of the gene ontologies, and the Hallmark GSEA pathways datasets^104,105^. Pathways were filtered out if they contained ≥ 400 or ≤ 10 genes. Pathways were considered to be significant if they had a Padj ≤ 0.01. GSEA plots were made using the Gseaplot2 function of ClusterprofileR. Differentially activated upstream regulators were assessed using Qiagen’s ingenuity pathway analysis (IPA) tool (QIAGEN, https://digitalinsights.qiagen.com/IPA). Upstream regulators that had an absolute z-score of < 0.2 or < 20% gene coverage were filtered out. Outputs of the Qiagen IPA platform were converted into a tab-delimited format, which was used to create bar plots using the R ggplot2 package.

### Statistical analyses

All analyses were performed using R Statistical Software (v4.1.1; R Core Team 2021). All bar plots represent mean values unless otherwise noted. Error bars represent 1.5 times the interquartile range between the 25th and 75th percentiles unless otherwise noted. Statistical significance between experimental groups was determined using a Wilcoxon rank-sum test and corrected with the Benjamini-Hochberg method unless otherwise noted in the figure legends. Differences were considered significant with P values < 0.05 (denoted by *), P < 0.01 (**), and P < 0.001 (***). The number of patients per experimental group is indicated in figure legends.

## Author Contributions

M.M. and S.F. designed experiments, M.M., Y.W., and W.L conducted experiments, M.M., and S.F. analyzed data and interpreted the results. M.M. and S.F. wrote the manuscript. All authors read and approved the final version of the manuscript.

## Conflicts of interest

S.F. is a scientific cofounder of CalciMedica; the other coauthors declare no conflict of interest.

## Funding Sources

This study was funded by National Institutes of Health (NIH) grant R01DE027981 (to R.S.L. and S.F.), NIH grant EY030917 (to S.C.), U01DE028891 (to Caroline H. Shiboski, SICCA), NIH F30 training grant AI164803 (to A.Y.T.), an award from the Colton Center for Autoimmunity at NYU (to S.F.), and Hunan Province Graduate Student Research and Innovation Project CX20190160 from Central South University, Changsha, Hunan, China (to W.L.).

## Data Availability

All raw sequencing data has been deposited in the Gene Expression Omnibus (GEO), accession number GSE253568.

---

**Table S1.**
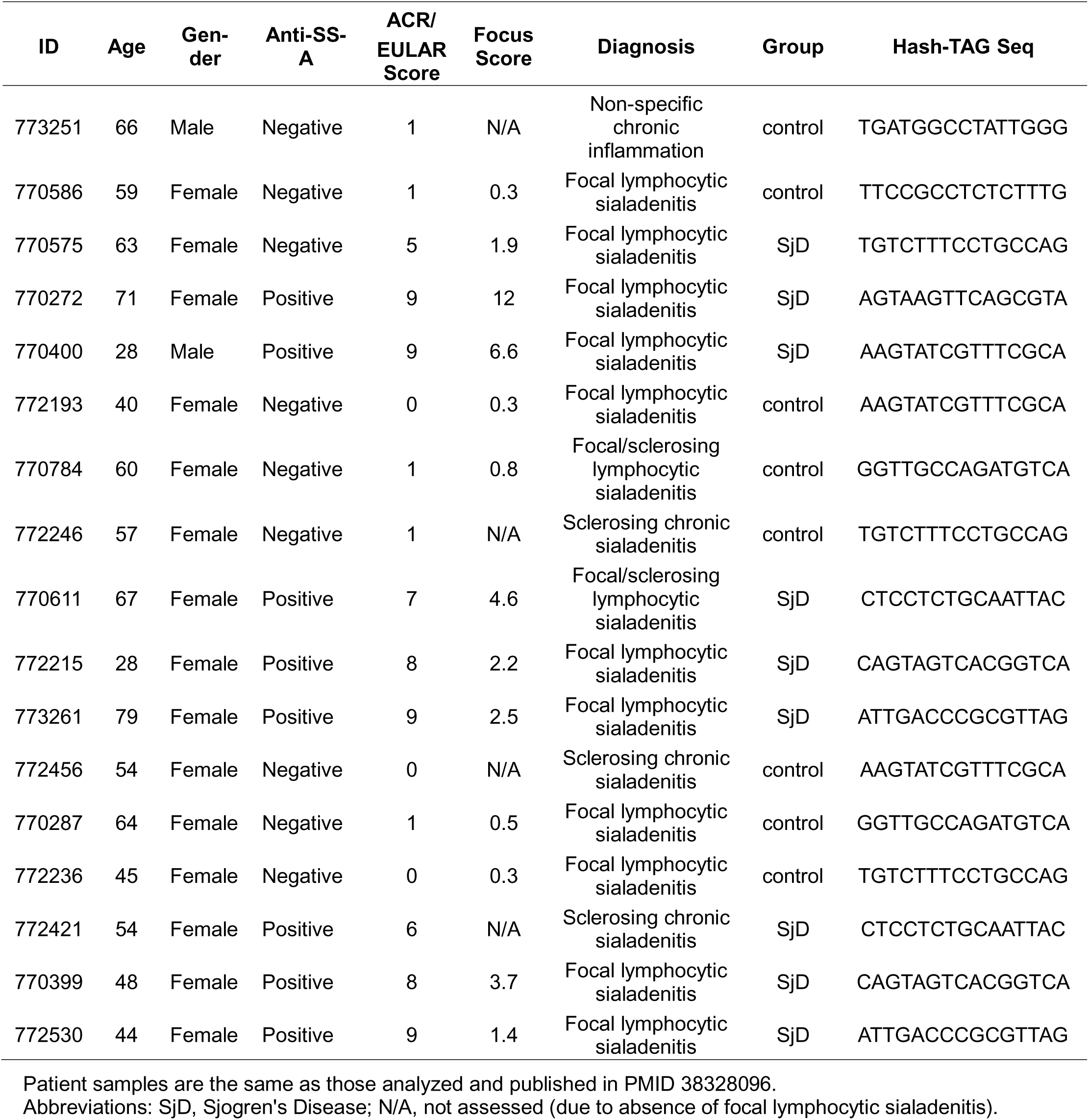
Patient information.

**Table S2.**
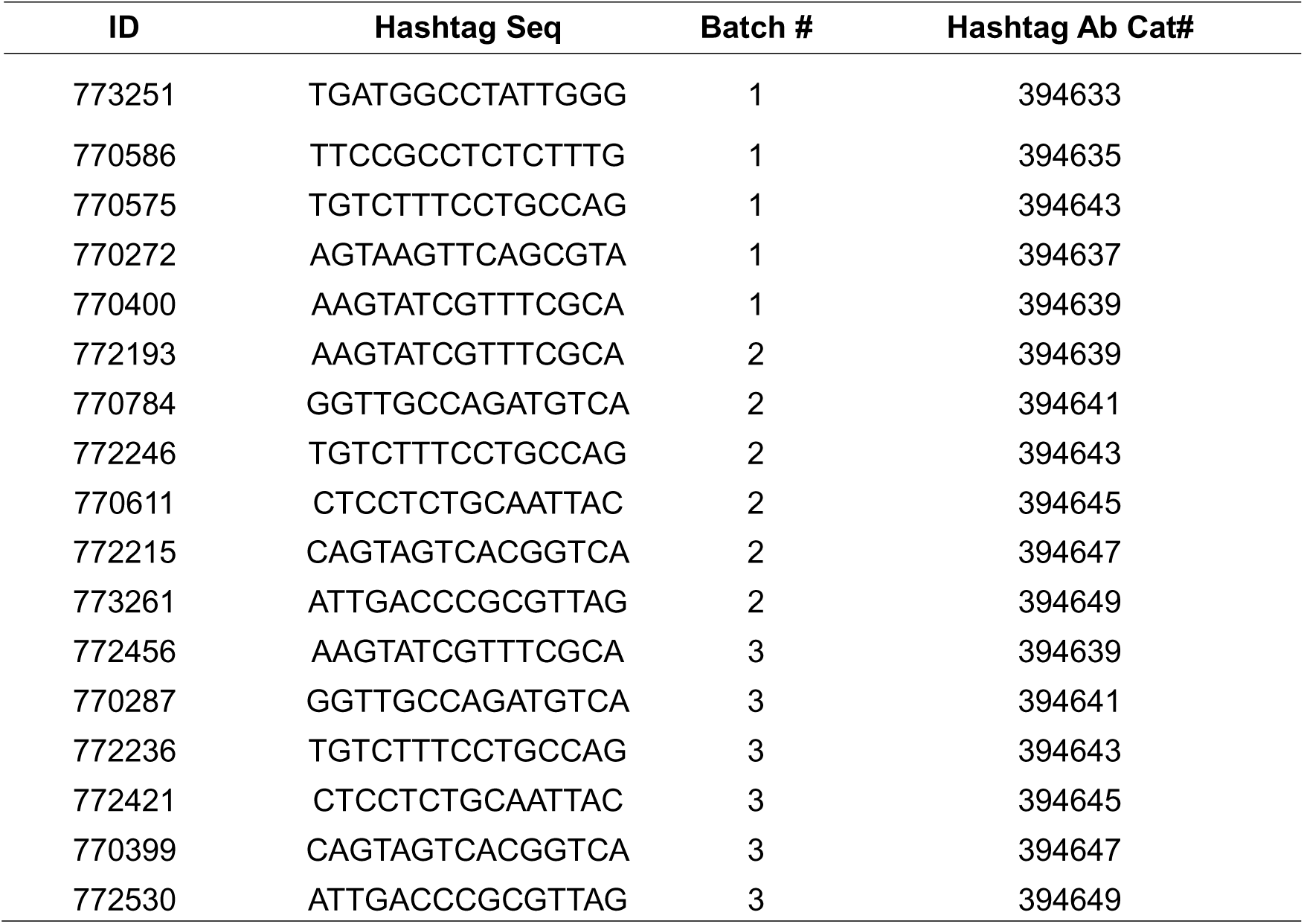
Hash tagging antibody information for each library.

## SUPPLEMENTARY FIGURE LEGENDS

**Figure S1.**
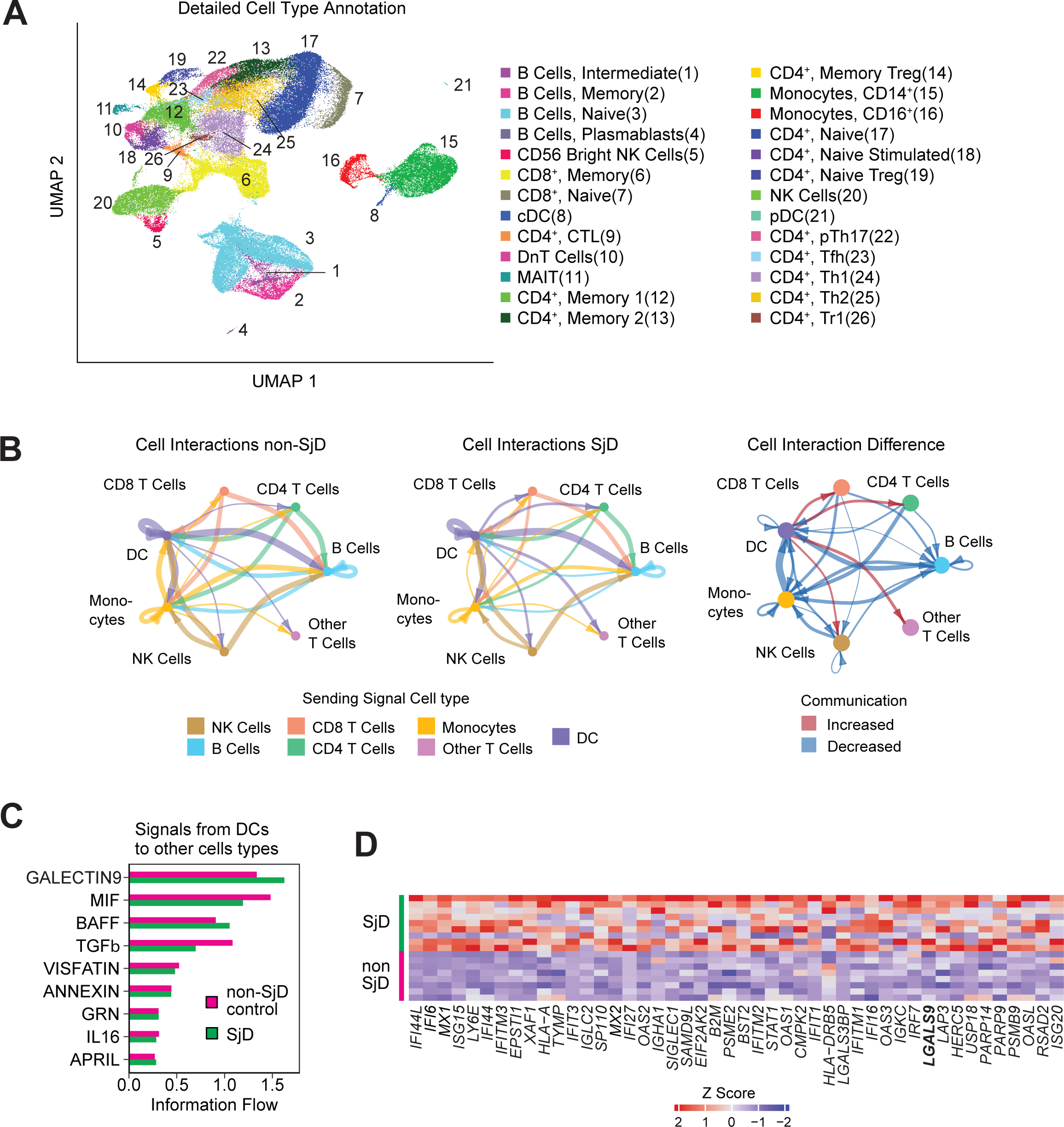
Detailed cell type annotation and analysis of cell-to-cell communication of SjD patients’ PBMC. **A)** Aggregate UMAP plot of 26 identified immune cell subsets within 105k peripheral blood mononuclear cells (PBMC) from 9 SjD patients and 8 non-SjD controls analyzed across 3 batches. **B)** Circle plots of cell-to-cell communication via ligand-receptor paired expression in PBMCs of non-SjD controls (left) and SjD patients (middle). The weighted difference in cell-to-cell communication between non-SjD controls and SjD patients is shown on the right. Red arrows indicate increased differential communication, blue arrows indicate decreased communication. The ligand-receptor (L-R) database used here is the secreted signaling category of cellchatDB. Pairs were calculated using the Trimean method, with the top and bottom 25% pairs being trimmed before calculating the weighted value. L-R Pairs were filtered if they were not expressed in at least 10 cells of a cell type and had a P > 0.01. **C)** Bar plot for the ligand receptor pairs that were found to be dysregulated which also originated from dendritic cells in either SjD patients or non SjD controls. **D)** Heatmap of all significant DEGs in DCs of SjD patients and non-SjD controls shown in Figure 1D. Colors indicate Z scores differentially expressed genes. Genes were considered significant if captured in > 10% of cells with an absolute log_2_ FC > 0.1 and Padj < 0.1.

**Figure S2.**
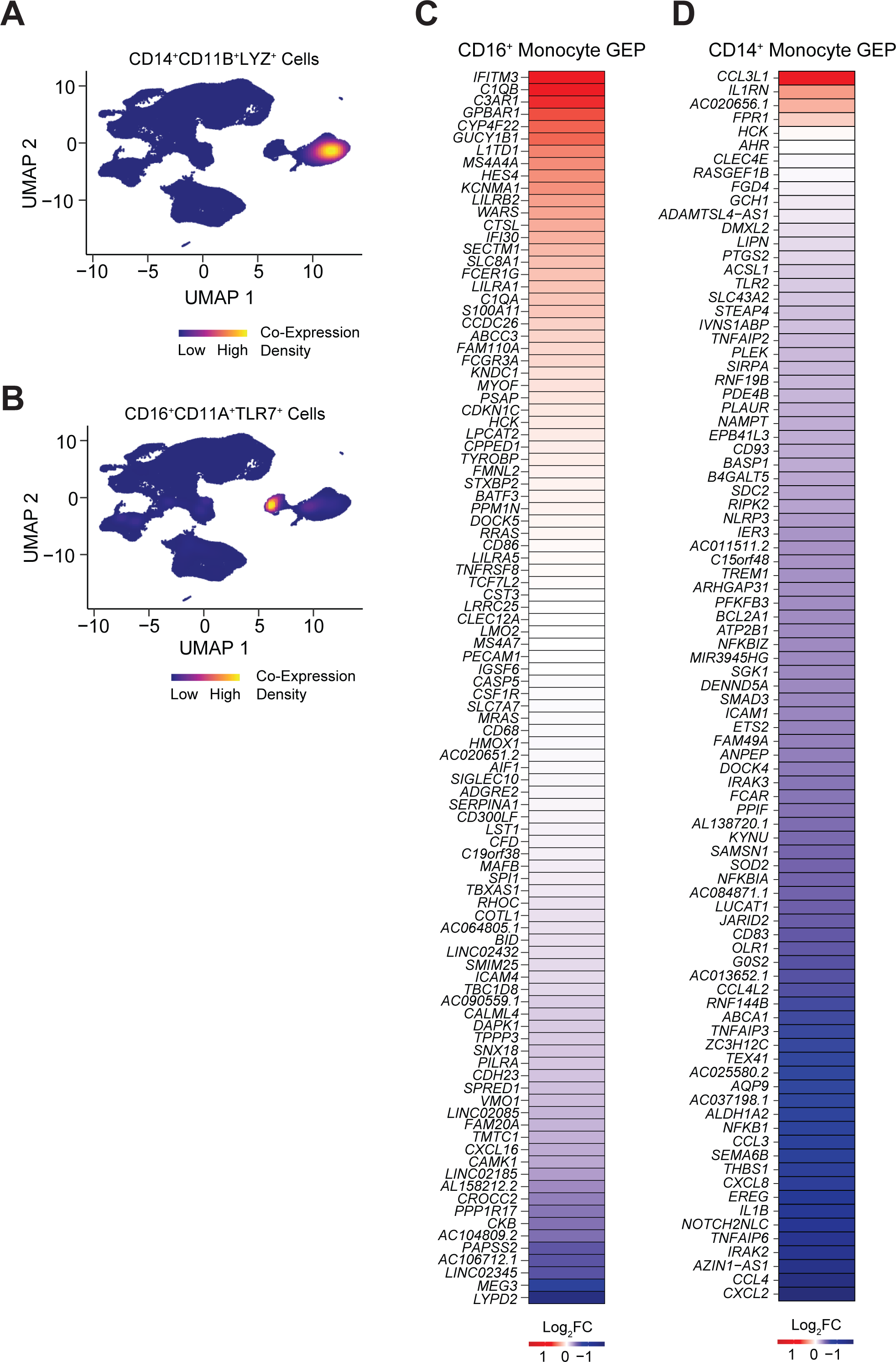
Characterization of CD14^+^ and CD16^+^ monocyte populations in SjD patients. **A)** Coexpression density plot of CD14^+^ monocyte marker genes overlayed onto the UMAP of all PBMCs. **B)** Coexpression density plot of CD16^+^ monocyte marker genes overlayed onto the UMAP of all PBMCs. **C)** Heatmap of DEGs contained in the *CD16^+^ monocyte* GEP in CD16^+^ monocytes of SjD patients and non-SjD controls. Colors depict log_2_ fold changes (FC) in expression. **D)** Heatmap of DEGs contained in the *CD14^+^ monocyte* GEP in CD14^+^ monocytes of SjD patients and non-SjD controls. Colors depict log_2_ fold changes (FC) in expression.

**Figure S3.**
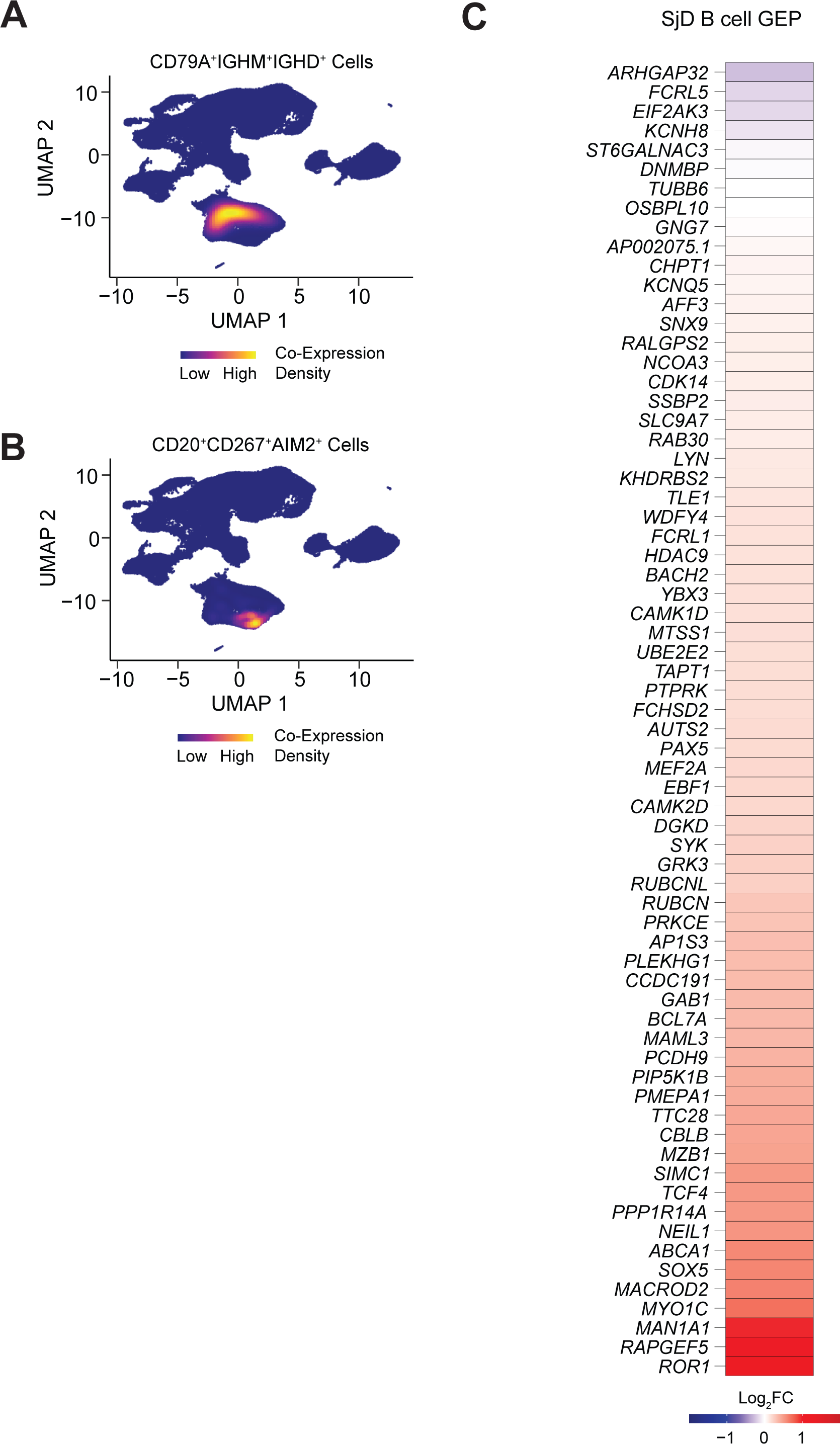
Characterization of B cell populations in SjD patients. **A)** Coexpression density plot of naïve B cell marker genes overlayed onto the UMAP of all PBMCs. **B)** Coexpression density plot of memory B cell marker genes overlayed onto the UMAP of all PBMCs. **C)** Tile plot of DEGs contained in the *SjD B cell* GEP in all B cells of SjD patients and non-SjD controls. Colors depict log_2_ fold changes (FC) in expression.

**Figure S4.**
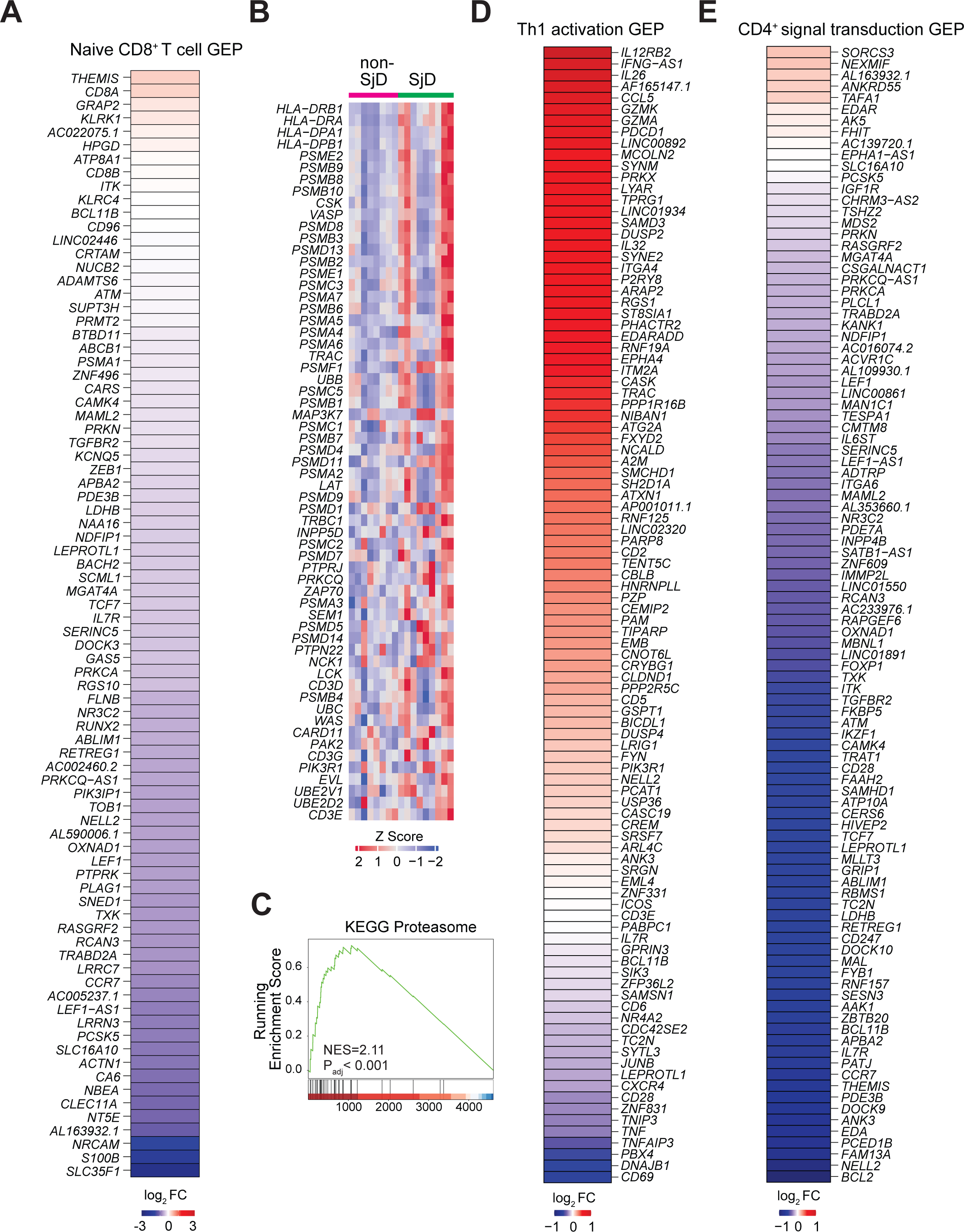
Characterization of CD8^+^ and CD4^+^ T cell populations in SjD patients. **A)** Heatmap of DEGs contained in the *naive CD8^+^ T cell* GEP in all CD8^+^ T cells of SjD patients and non-SjD controls. Colors depict log_2_ fold changes (FC) in expression. **B)** Heatmap of DEGs in CD4^+^ T cells of SjD patients and non-SjD controls identified using the Reactome TCR signaling gene set. **C)** GSEA of DEGs in all CD4^+^ T cells of SjD patients and controls. Shown is an enriched gene set related to proteosome-related genes. **D)** Heatmap of DEGs contained in the *Th1 activation* GEP in all CD4^+^ T cells of SjD patients and non-SjD controls. Colors depict log_2_ fold changes (FC) in expression. **E)** Heatmap of DEGs contained in the *CD4^+^ T cell signal transduction* GEP in all CD4^+^ T cells of SjD patients and non-SjD controls. Colors depict log_2_ fold changes (FC) in expression. Z scores shown in (B) were calculated using the normalized average expression per cell for each gene across all samples.

